# Zika virus replicates in the vagina of mice with intact interferon signaling

**DOI:** 10.1101/2022.02.21.481392

**Authors:** Cesar A. Lopez, Sarah J. Dulson, Helen M. Lazear

## Abstract

Zika virus (ZIKV) is unusual among flaviviruses in its ability to spread between humans through sexual contact, as well as by mosquitoes. Sexual transmission has the potential to change the epidemiology and geographic range of ZIKV compared to mosquito-borne transmission and potentially could produce distinct clinical manifestations, so it is important to understand the host mechanisms that control susceptibility to sexually transmitted ZIKV. ZIKV replicates poorly in wild-type mice following subcutaneous inoculation, so most ZIKV pathogenesis studies use mice lacking IFN-αβ signaling (e.g. *Ifnar1*^-/-^). However, we found that wild-type mice support ZIKV replication following intravaginal infection, although the infection remained localized to the lower female reproductive tract. Vaginal replication was not a unique property of ZIKV, as other flaviviruses that generally are restricted in wild-type mice also were able to replicate in the vagina. Vaginal ZIKV infection required a high-progesterone state (pregnancy or pre-treatment with depot medroxyprogesterone acetate (DMPA)), identifying a key role for hormonal status in susceptibility to vaginal infection. Progesterone-mediated susceptibility did not appear to result from a compromised epithelial barrier, blunted antiviral gene induction, or changes in vaginal leukocyte populations, leaving open the mechanism by which progesterone confers susceptibility to vaginal ZIKV infection. Progesterone treatment is a key component of mouse vaginal infection models for herpes simplex virus and *Chlamydia*, but the mechanisms by which DMPA increases susceptibility to those pathogens also remain poorly defined. Understanding how progesterone mediates susceptibility to ZIKV vaginal infection may provide insights into host mechanisms influencing susceptibility to diverse sexually transmitted pathogens.

**IMPORTANCE:** Zika virus (ZIKV) is transmitted by mosquitoes, similarly to other flaviviruses. However, ZIKV is unusual in its ability also to spread through sexual transmission. We found that ZIKV was able to replicate in the vaginas of wild-type mice, even though these mice do not support ZIKV replication by other routes, suggesting that the vagina is particularly susceptible to ZIKV infection. Vaginal susceptibility was dependent on a high progesterone state, which is a common feature of mouse vaginal infection models for other pathogens, through mechanisms that have remained poorly defined. Understanding how progesterone mediates susceptibility to ZIKV vaginal infection may provide insights into host mechanisms that influence susceptibility to diverse sexually transmitted pathogens.

## INTRODUCTION

The unprecedented size of the 2015-2016 Zika virus pandemic in the Americas, in which millions of people were infected, revealed new disease manifestations and transmission mechanisms, including congenital infection and sexual transmission (1). Flaviviruses are transmitted to humans by arthropod vectors (mosquitoes and ticks), and ZIKV is the first example of a flavivirus that spreads between humans via sexual transmission (2). The first report of ZIKV sexual transmission pre-dates the 2015-2016 epidemic and resulted from ZIKV infection in Africa (3), suggesting that sexual transmission is a general property of ZIKV, rather than a new trait coincident with its emergence in the Americas. The ability of ZIKV to spread via sexual transmission in addition to mosquito-borne transmission expands the geographic range over which ZIKV transmission can occur, could change the epidemiology of ZIKV even in areas with mosquito-borne transmission, and has the potential to produce distinct pathologic outcomes if congenital infection occurs via an ascending route rather than a hematogenous transplacentalroute. Thus, it is important to understand the antiviral mechanisms that ZIKV may encounter in the vagina that are distinct from antiviral mechanisms present at the skin following mosquito inoculation.

Mouse models of ZIKV vaginal infection involve pre-treating mice with progesterone, based on well-established infection models for herpes simplex virus (HSV) and *Chlamydia muridarum* (4, 5). The mechanism by which progesterone makes mice susceptible to ZIKV remains unknown but has been hypothesized to be due to a combination of thinned epithelium, infiltrating immune cells susceptible to ZIKV infection, or deficiencies in antiviral signaling due to decreased expression of antiviral sensing genes (6-8). ZIKV replication is restricted by the type I interferon (IFN-αβ) response in mice because ZIKV is unable to antagonize mouse STAT2 (9, 10). Thus, mouse models of ZIKV pathogenesis, including those investigating vaginal infection, typically use mice deficient in IFN-αβ signaling, usually through genetic loss of the IFN-αβ receptor (*Ifnar1*^*-/-*^) alone or in combination with the IFN-γ receptor, or by treatment of wild-type mice with an IFNAR1-blocking monoclonal antibody (7, 11-15).

Here we show that although wild-type mice largely are resistant to ZIKV infection via footpad inoculation, vaginal inoculation results in productive local ZIKV replication. We further show that permissiveness to vaginal ZIKV replication is regulated by progesterone, in a manner dominant to IFN-αβ signaling, identifying a key role for hormonal status in susceptibility to vaginal infection. Vaginal replication was not a unique property of ZIKV, as other flaviviruses that generally are restricted in wild-type mice also were able to replicate in the vagina. Progesterone-mediated susceptibility did not appear to result from a compromised epithelial barrier, blunted antiviral gene induction, or changes in vaginal leukocyte populations, leaving open the mechanism by which progesterone confers susceptibility to vaginal ZIKV infection.

## RESULTS

### Wild-type mice support ZIKV replication after intravaginal inoculation

Mouse models of ZIKV pathogenesis typically employ mice lacking IFN-αβ signaling (e.g. *Ifnar1*^*-/-*^) to achieve robust infection, as wild-type mice sustain only minimal replication following subcutaneous inoculation (11, 16, 17). Accordingly, in seeking to define host mechanisms that control ZIKV infection in the female reproductive tract, we compared ZIKV replication in wild-type and *Ifnar1*^-/-^ mice following intravaginal inoculation. We pre-treated wild-type and *Ifnar1*^*-/-*^ mice with depot medroxyprogesterone acetate (DMPA) (a standard component of mouse vaginal infection models for herpes simplex virus (HSV), *Chlamydia*, and ZIKV), then 5 days later infected with 1000 FFU of ZIKV via intravaginal instillation (Figure 1). We assessed viral replication in the vagina by collecting vaginal washes 2, 4, 6, and 8 days post-infection (dpi) and measuring ZIKV RNA by qRT-PCR. We found that viral loads increased from 2 through 8 dpi, indicating productive replication in the vagina. In contrast to minimal viral replication observed in wild-type mice after subcutaneous inoculation in the footpad (11, 16, 17), we observed similar ZIKV replication kinetics and RNA burden in the vaginas of wild-type compared to *Ifnar1*^*-/-*^ mice, with the only significant difference being higher viral loads in wild-type mice at 2 dpi (Figure 1A). Although wild-type mice supported ZIKV replication in the vagina, they did not support systemic infection as viremia was detected only in *Ifnar1*^-/-^ mice (Figure 1B). Likewise, *Ifnar1*^*-/-*^ mice supported ascending infection into the upper female reproductive tract (uterus, ovary, and oviduct) whereas ZIKV infection in wild-type mice was restricted to the lower female reproductive tract (cervix) (Figure 1C). To confirm that the ZIKV RNA we detected in vaginal washes represented replicating virus, we inoculated wild-type mice with either infectious ZIKV or UV-inactivated virus and measured viral RNA in vaginal washes collected 2 through 8 dpi. No ZIKV RNA was detected in vaginal washes from mice inoculated with UV-inactivated virus, further supporting that the viral RNA detected in vaginal washes results from productive infection (Figure 1D). Altogether, these results show that ZIKV can replicate in the vagina of wild-type mice, but that IFN-αβ signaling restricts systemic spread.

**Figure 1:**
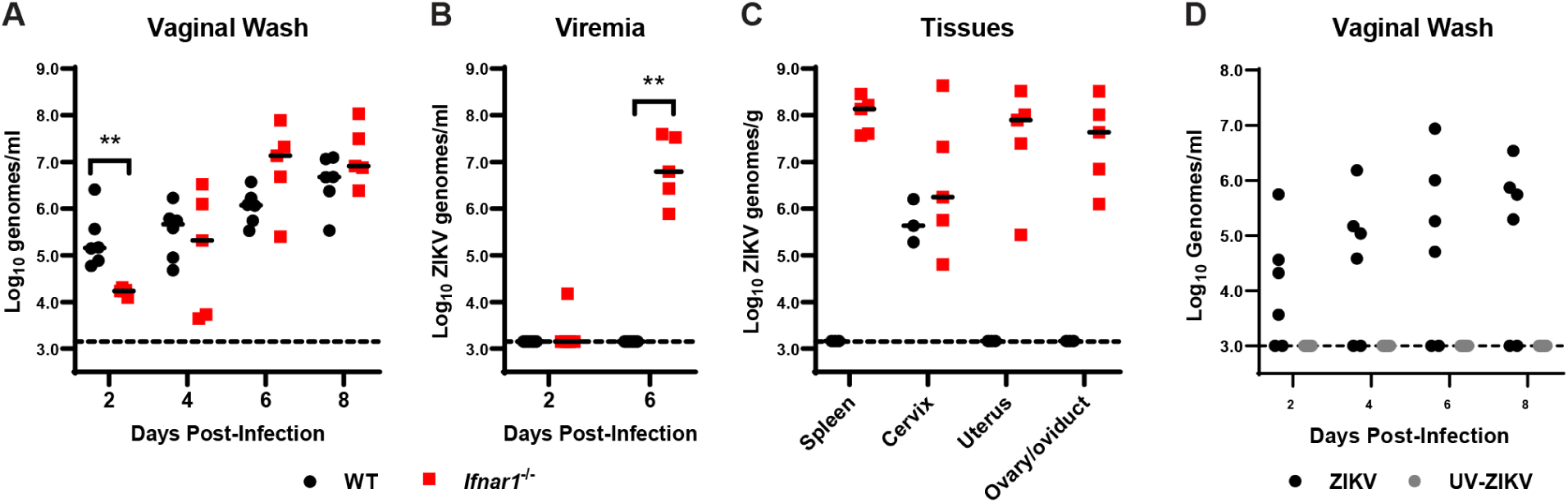
WT mice are susceptible to ZIKV vaginal infection. 6 to 7 week-old mice were pre-treated with 2 mg of DMPA and inoculated with 1000 FFU of ZIKV by intravaginal instillation 5 days later. **A-C**. Viral RNA extracted from vaginal washes (**A**), serum (**B**), or tissues (**C**) of wild-type and *Ifnar1*^-/-^ mice was measured by qRT-PCR. Data represent 5-6 (**A-B**) or 3-5 (**C**) mice per group combined from 2 independent experiments. WT and *Ifnar1*^-/-^ groups were compared by Mann-Whitney test with adjustment for multiple comparisons (*, P <0.05; **, P <0.01). **D**. WT mice were inoculated intravaginally with 1000 FFU of mock-inactivated or UV-inactivated ZIKV. Viral RNA was extracted from vaginal washes and measured by qRT-PCR. Data represent 6 mice per group combined from 2 independent experiments.

In addition to the antiviral effects of IFN-αβ, type III IFNs (IFN-λ) contribute to antiviral immunity at epithelial barriers (18). IFN-λ has been reported to restrict HSV infection in the vagina (19) and to restrict ZIKV infection in the vagina when IFN-αβ signaling is inhibited by administration of an IFNAR1-blocking antibody (15). To test whether IFN-λ controls vaginal ZIKV infection in mice with intact IFN-αβ signaling, we used mice lacking the IFN-λ receptor (*Ifnlr1*^-/-^). We treated *Ifnlr1*^+/-^ and *Ifnlr1*^-/-^ mice with DMPA and infected with 1000 FFU of ZIKV by intravaginal instillation. We measured viral loads in vaginal washes by qRT-PCR and found no significant difference between *Ifnlr1*^+/-^ and *Ifnlr1*^-/-^ mice, suggesting that IFN-λ signaling does not restrict ZIKV replication in the vagina in this model (Figure 2).

**Figure 2:**
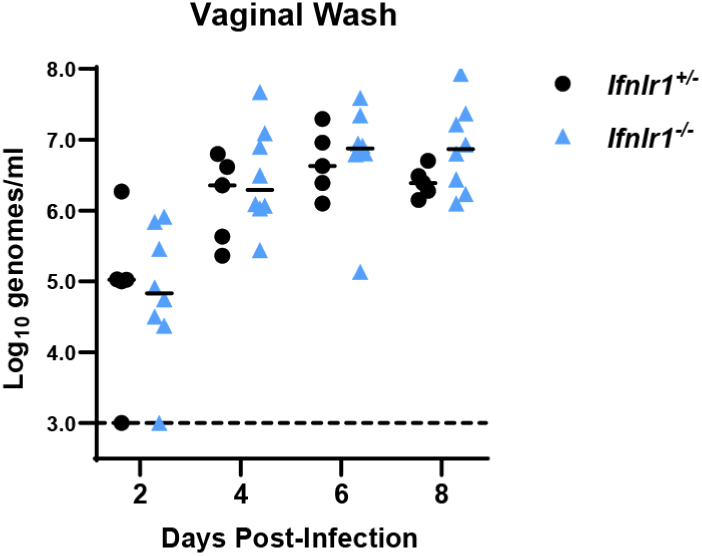
IFN-λ does not restrict ZIKV infection in the vagina. 5-6 week-old mice lacking (*Ifnlr1*^-/-^) or retaining (*Ifnlr1*^+/-^) IFN-λ signaling were pre-treated with 2 mg of DMPA and inoculated 5 days later with 1000 FFU of ZIKV by intravaginal instillation. Viral RNA was measured from vaginal washes by qRT-PCR. *Ifnlr1*^-/-^ and *Ifnlr1*^+/-^ groups were compared by Mann-Whitney with adjustment for multiple comparisons. Data are combined from 3 independent experiments.

### A high-progesterone state is required for vaginal ZIKV infection

Pre-treatment with DMPA is a standard component of mouse models of vaginal infection with diverse pathogens including HSV, *Chlamydia*, and ZIKV (4-7). Since we found that wild-type mice were susceptible to ZIKV infection via an intravaginal but not a subcutaneous inoculation route, we considered whether DMPA treatment rendered mice susceptible to systemic ZIKV infection. We treated wild-type mice with DMPA or PBS, then 5 days later infected with 1000 FFU of ZIKV via intravaginal instillation or subcutaneous inoculation in the footpad and measured viral RNA in vaginal wash and in serum by qRT-PCR. As expected, DMPA treatment increased the permissiveness of wild-type mice to intravaginal infection: ZIKV RNA was detected in the vaginal wash from 10 of 10 DMPA-treated mice compared to only 5 of 10 PBS-treated mice (3 of which were positive on only a single day) and DMPA-treated mice sustained higher viral loads in the vagina than PBS-treated mice (Figure 3A). Consistent with previous experiments, DMPA-treated wild-type mice supported ZIKV replication in the vagina but no ZIKV RNA was detected in the serum following intravaginal inoculation (Figure 3B). Furthermore, no ZIKV RNA was detected in the serum of mice inoculated by footpad regardless of DMPA treatment (Figure 3B), indicating that DMPA treatment was not sufficient to render wild-type mice broadly susceptible to ZIKV infection. Although *Ifnar1*^*-/-*^ mice are highly susceptible to ZIKV infection by subcutaneous inoculation, productive vaginal infection required DMPA treatment (1 of 10 PBS-treated mice infected compared to 9 of 9 DMPA-treated) (Figure 3C); all *Ifnar1*^-/-^ mice with productive vaginal infection subsequently developed viremia (Figure 3D). These results demonstrate a key role for progesterone in susceptibility to vaginal ZIKV infection, even in the context of immunodeficient mice that are otherwise highly susceptible to ZIKV infection.

**Figure 3:**
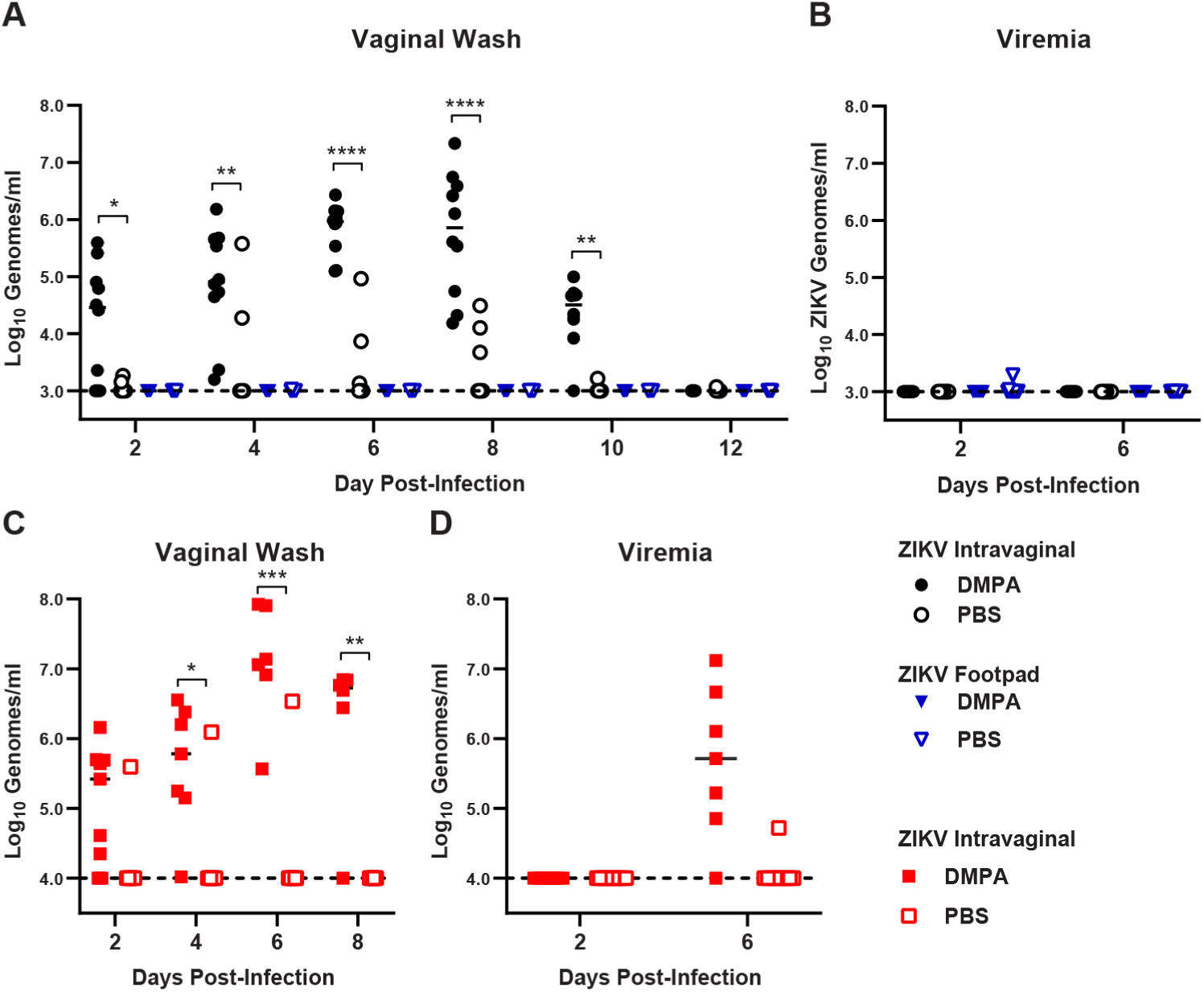
DMPA does not sensitize WT mice to ZIKV infection by footpad inoculation. 6-week-old wild-type (**A-B**) or *Ifnar1*^-/-^ mice (**C-D**) were pre-treated with either PBS or 2 mg of DMPA then infected with 1000 FFU of ZIKV by intravaginal instillation or subcutaneous inoculation in the footpad. Viral RNA in vaginal washes (**A** and **C**) or serum (**B** and **D**) was measured by qRT-PCR. Data represent 9 or 10 mice per group combined from 2 independent experiments. PBS and DMPA treated groups were compared by two-way ANOVA with multiple comparison correction (*, P <0.05; **, P <0.01; ***, P < 0.001; ****, P <0.0001).

Since congenital infection is an important manifestation of ZIKV infection, and pregnancy is a high-progesterone state (20), we evaluated vaginal ZIKV infection in pregnant mice (without DMPA treatment). We mated 7-to-10-week old wild-type dams with wild-type sires and inoculated 7 days post-mating (roughly one-third of gestation) intravaginally with 1000 FFU of ZIKV. We collected vaginal washes and serum and measured ZIKV RNA by qRT-PCR to assess local replication in the vagina and systemic spread, and all mice were harvested at 8 dpi to assess congenital infection. Pregnant mice supported vaginal ZIKV replication (viral RNA detected in the vaginal wash from 4 of 5 pregnant mice) but ZIKV RNA was not detected in the vaginal lavage of non-pregnant mice (0 of 12 mice) (Figure 4A). Consistent with our observations in non-pregnant wild-type mice, pregnant wild-type mice did not support systemic ZIKV spread, as ZIKV RNA was not detected in serum, even in the context of robust replication in the vagina (Figure 4B).

**Figure 4:**
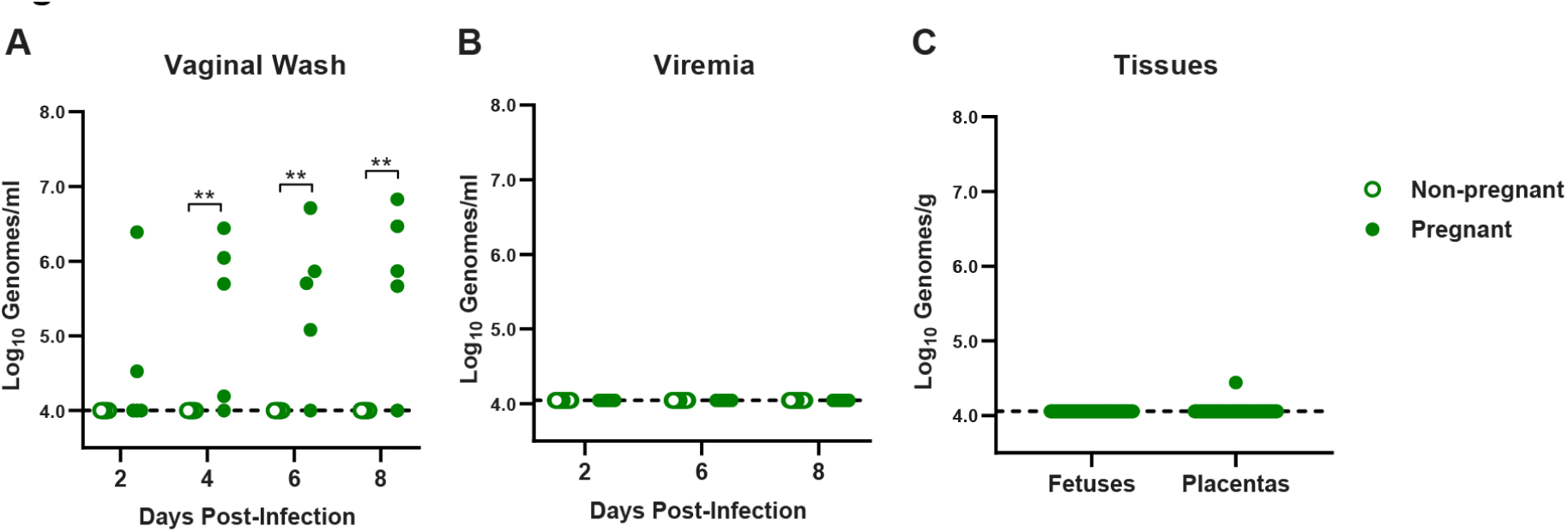
Pregnant WT mice are susceptible to intravaginal ZIKV infection. 7-to-10 week-old wild-type dams were mated with WT sires and inoculated 7 days afterwards intravaginally with 1000 FFU of ZIKV. Viral RNA was measured by qRT-PCR in vaginal washes (**A**), serum (**B**), or fetal tissues harvested at day 8 post-infection (**C**). Data are combined from 5 pregnant and 12 non-pregnant dams and 40 placentas and fetuses from 2 independent experiments. Pregnant and non-pregnant groups were compared by Mann-Whitney, adjusted for multiple comparisons (**, P <0.01).

Additionally, ZIKV RNA was detected in only 1 of the 40 placentas and none of the corresponding fetuses (Figure 4C), consistent with the lack of ascending or systemic infection we observed after vaginal ZIKV inoculation in DMPA-treated non-pregnant wild-type mice (Figure 1B). Altogether these data suggest that a high progesterone state (DMPA treatment or pregnancy) is required for vaginal permissiveness to ZIKV infection, and that vaginal infection is not sufficient for maternal-fetal transmission.

### The vagina is permissive to replication of diverse IFN-αβ-restricted flaviviruses

ZIKV is unique among flaviviruses in its ability to spread among humans via both vector-borne (mosquito) and vector-independent (sexual) transmission routes. To assess whether this reflects an unusual vaginal tropism of ZIKV, we evaluated vaginal infection with 3 additional flaviviruses, Spondweni virus (SPOV), Usutu virus (USUV), and dengue virus (DENV). These flaviviruses were selected because, like ZIKV, they replicate poorly in wild-type mice following subcutaneous inoculation (21-23). Wild-type mice were treated with DMPA 5 days prior to intravaginal inoculation with 1000 FFU of ZIKV, SPOV, USUV, or 10,000 FFU of DENV3 and viral RNA was measured by qRT-PCR from vaginal washes 2, 4, 6, and 8 dpi (Figure 5). Viral RNA was detected in vaginal washes after ZIKV, USUV, and SPOV infection, suggesting that these viruses could replicate in the vagina of wild-type mice and at levels similar to ZIKV. In contrast,

**Figure 5:**
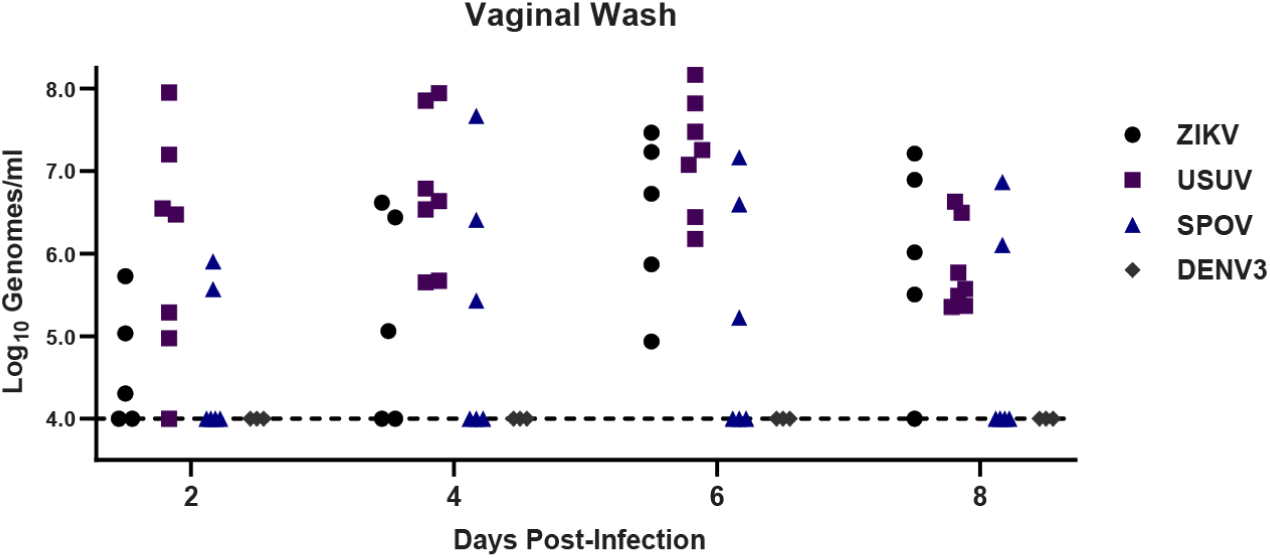
Diverse flaviviruses replicate in the vagina of WT mice. 6-week-old wild-type mice pre-treated with 2 mg of DMPA were inoculated with 1000 FFU of ZIKV, Usutu virus (USUV), Spondweni virus (SPOV), or dengue virus (DENV3) by intravaginal instillation. Viral RNA was measured from vaginal washes by qRT-PCR.

DENV3 RNA was not detected. To test whether the vagina is permissive to other RNA viruses that generally are restricted by innate antiviral responses in wild-type mice (24, 25), we inoculated wild-type mice intravaginally with 1000 FFU of rubella virus (*Matonaviridae*) or 5 × 10^8^ genome equivalents of hepatitis A virus (*Picornaviridae*) but detected no viral RNA in vaginal washes at any of the time points evaluated through 8 dpi (data not shown). Altogether, these data show that vaginal infection is not a unique property of ZIKV among flaviviruses. Rather, in wild-type mice the vagina is more permissive to flavivirus replication compared to other inoculation sites but does not allow unrestricted replication of all RNA viruses.

### ZIKV infection in the vagina is localized to the epithelium

To better define the location of the cells targeted by ZIKV in the vagina, we treated wild-type mice with DMPA, infected them intravaginally, and detected ZIKV RNA in vaginal tissue using RNAscope *in situ* hybridization (Figure 6). ZIKV positive cells were infrequent and sporadically distributed in the vagina, but they tended to be clusters of adjacent epithelial cells located along the vaginal lumen, with little staining in the parenchyma. We detected ZIKV staining in 0 of 3 mice at 2 dpi, 1 of 2 at 4 dpi, 3 of 3 at 6 dpi, 2 of 2 at 8 dpi, and 0 of 5 at 10 dpi. The largest clusters of infected cells were detected at 6 dpi. There was no tendency for infected cells to be nearer to the cervix or nearer to the vaginal opening. No sections from infected mice exhibited leukocyte infiltrate into to the vaginal tissue relative to uninfected DMPA-treated mice. Altogether, these results indicate that ZIKV infection in the vagina primarily targets epithelial cells, rather than the leukocytes that are the main targets of ZIKV systemic infection (26, 27), and that infected cells are not associated with a pronounced immune infiltrate.

**Figure 6:**
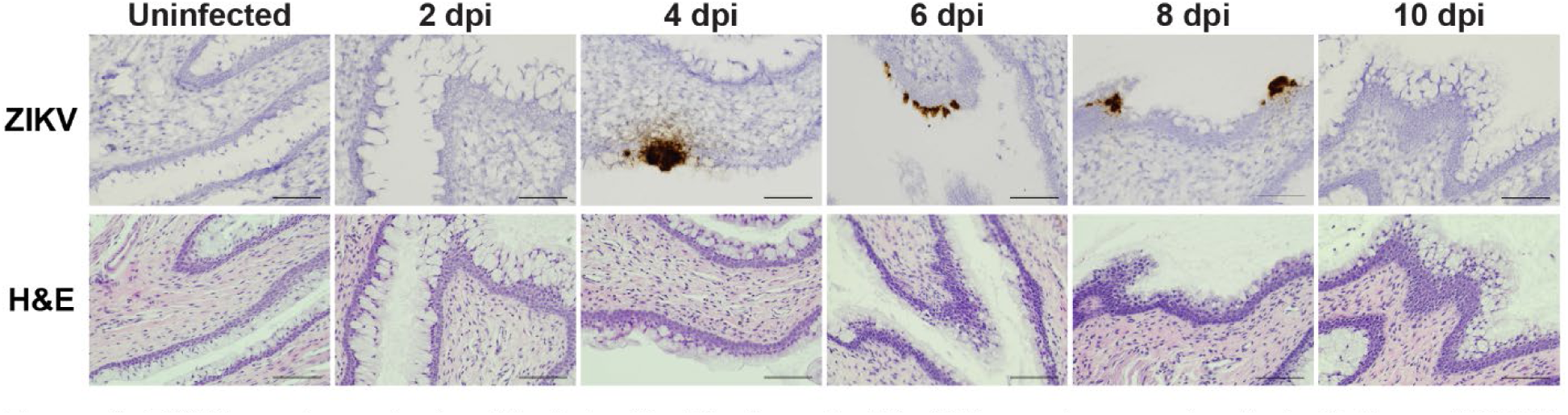
ZIKV targets vaginal epithelial cells. 5 to 6 week-old wild-type mice were treated with 2 mg of DMPA and 5 days later infected with 1000 FFU ZIKV intravaginally. Vaginal tissue was harvested 2 to 10 dpi, paraffin embedded, and adjacent sections were stained for ZIKV RNA or H&E. Each image is a single field at 20x (scale bar: 100 µm).

### A physically compromised vaginal epithelial barrier is not sufficient to render wild-type mice susceptible to ZIKV infection

DMPA treatment induces a diestrus-like state in mice, including a vaginal epithelium that is thinned and lacks extensive keratinization (7). Since we found that ZIKV infects epithelial cells in the vagina, we hypothesized that a thinned epithelial barrier is more easily targeted by ZIKV, explaining the requirement of DMPA for susceptibility of wild-type and *Ifnar1*^*-/-*^ mice to vaginal ZIKV infection. To test whether an impaired epithelial barrier could overcome the requirement for DMPA treatment, we abraded the vaginal epithelium of wild-type mice with an interdental brush prior to intravaginal inoculation with 1000 FFU of ZIKV and measured ZIKV RNA in vaginal washes by qRT-PCR. However, vaginal infection was only detected in mice that were treated with DMPA, regardless of vaginal abrasion (Figure 7A) suggesting that a disrupted epithelial barrier is not sufficient for productive ZIKV infection in the vagina. Vaginal abrasion also did not facilitate ZIKV dissemination as ZIKV RNA was not detected in serum even from abraded mice (Figure 7B). In DMPA-treated mice, abrasion did not result in higher viral loads in vaginal washes, altogether suggesting that compromised epithelial barrier integrity is not the mechanism by which DMPA treatment promotes vaginal ZIKV infection.

**Figure 7:**
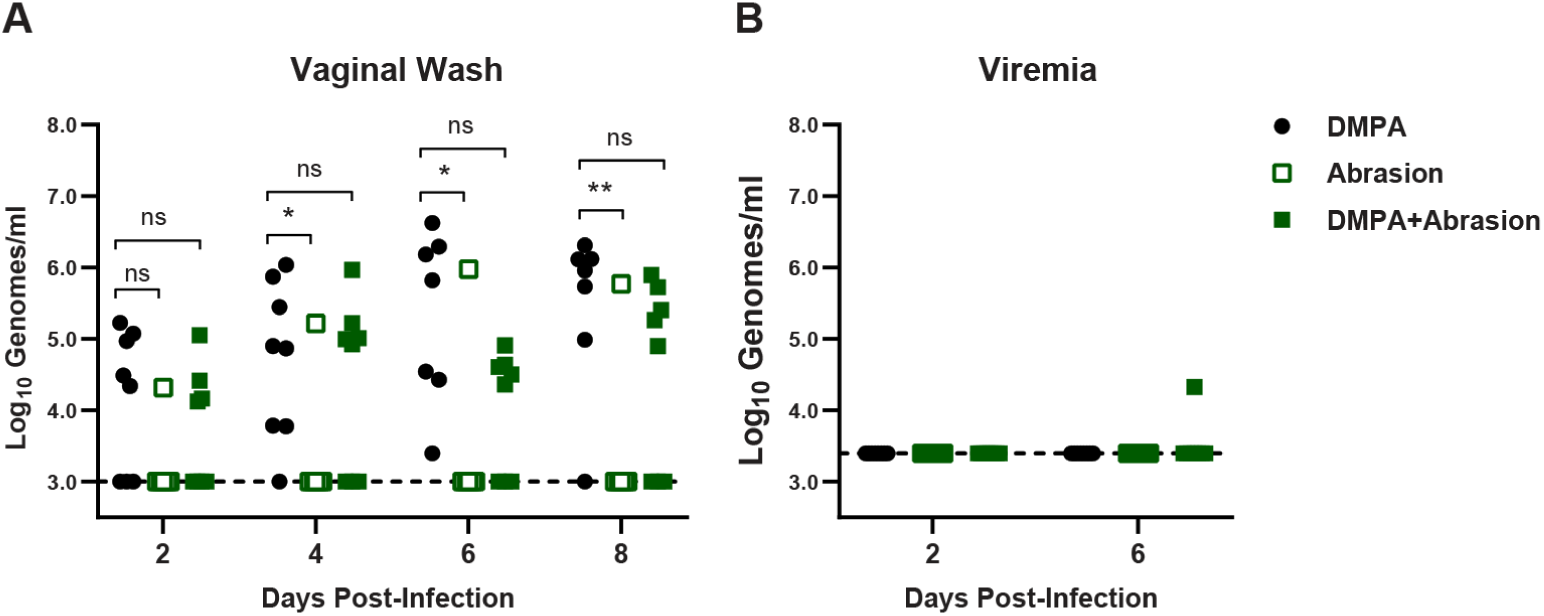
Vaginal abrasion is not sufficient to sensitize WT mice to ZIKV intravaginal infection. 6 week old wild-type mice were treated with 2 mg of DMPA 5 days prior to inoculation, or vaginally abraded with an interdental brush immediately prior to inoculation with 1000 FFU ZIKV via vaginal instillation. Viral RNA in vaginal washes (**A**) or serum (**B**) was measured by qRT-PCR. Data represent 8 mice per group combined from 2 independent experiments. Abraded groups were compared to DMPA-only by two-way ANOVA, corrected for multiple comparisons (ns, not significant P >0.05; *, P <0.05; **, P <0.01).

### DMPA treatment does not diminish ISG expression or induction

We next considered whether DMPA treatment might inhibit the basal expression or induction of IFN-stimulated genes (ISGs) in the vagina, thereby permitting ZIKV replication. To test the effect of DMPA treatment on vaginal ISG expression, we treated wild-type mice with DMPA or PBS then 4 days later administered 50 µg of poly(I:C) intravaginally or intraperitoneally. One day after poly(I:C) treatment, we harvested tissues and measured expression of the canonical ISG *Ifit1* by qRT-PCR. We did not observe any DMPA-dependent change in *Ifit1* induction in the spleen or lower female reproductive tract (LFRT, vagina and cervix) following intravaginal or intraperitoneal poly(I:C) treatment (Figure 8A and B). In the upper female reproductive tract (UFRT, uterus and oviduct), DMPA increased *Ifit1* expression at baseline and induction in response to intravaginal poly(I:C) treatment (Figure 8C). DMPA potentially could selectively regulate expression of some ISGs but not *Ifit1*, or do so in response to viral infection but not poly(I:C) treatment, but our results do not support a model where DMPA-induced susceptibility to vaginal ZIKV infection is due to a broad inhibition of basal or induced ISG expression.

**Figure 8:**
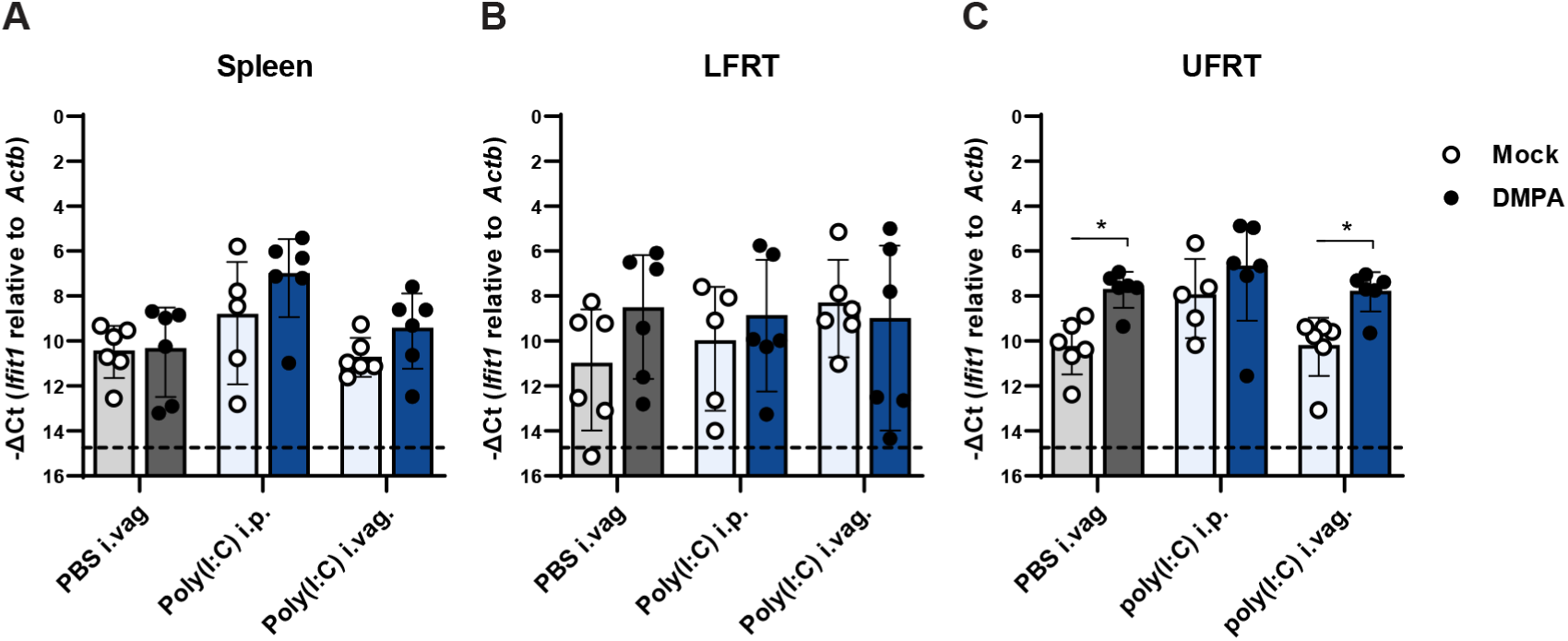
DMPA does not inhibit ISG expression. 5 to 6 week-old wild-type mice were treated with 2 mg of DMPA or PBS (mock). Four days later, mice were treated with 50 µg of poly(I:C) intravaginally (i.vag). or intraperitoneally (i.p.) or PBS i.vag. and tissues were harvested the following day. RNA was extracted from spleen (**A**), lower female reproductive tract (LFRT, vagina and cervix) (**B**), or upper female reproductive tract (UFRT, uterus and oviduct) (**C**). *Ifit1* expression was measured as -ΔCt normalized to *Actb*. Mock and DMPA-treated groups were compared by Mann-Whitney with adjustment for multiple comparisons (*, P <0.05).

### DMPA treatment does not change vaginal or systemic leukocyte populations

We next considered whether DMPA might alter leukocyte populations in the vagina or systemically, which could facilitate ZIKV infection by suppressing antiviral immunity or by recruiting susceptible target cells to the site of infection. Wild-type mice were treated with DMPA alone, treated with DMPA then infected with ZIKV intravaginally 5 days later, or left untreated. Six days after ZIKV infection (11 days after DMPA treatment), cells were isolated from spleen, iliac lymph node, and LFRT. Total cell counts were calculated for T cells, NK cells, B cells, dendritic cells (DCs), eosinophils, monocytes, and neutrophils (markers and gating as defined in methods). ZIKV infection caused an increase in the number of B cells in the spleen and iLN and T cells in the spleen (Figure 9A-B), and an increase in the number of DCs in the LFRT (Figure 9C) compared to DMPA treatment alone. However, DMPA treatment alone caused no change in leukocyte populations in any of the tissues analyzed compared to untreated mice. Although DMPA potentially could affect specific leukocyte subsets not analyzed here, or affect activation states independently of cell numbers, our results suggest that DMPA-induced susceptibility to vaginal ZIKV infection does not result from a dramatic change in the immune milieu of the vagina.

**Figure 9:**
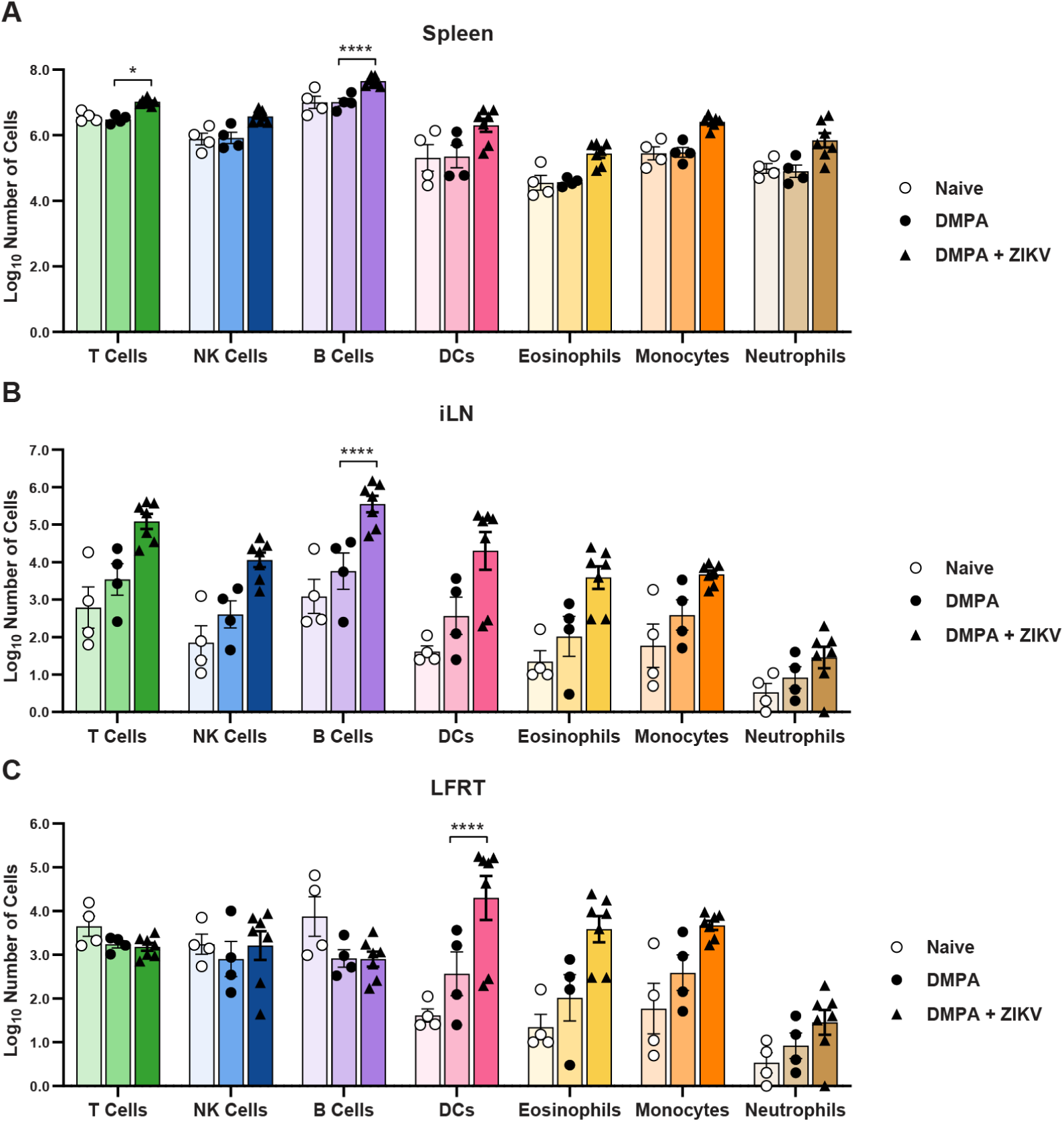
DMPA treatment alone does not impact systemic or vaginal leukocyte populations. 5 to 6 week-old wild-type mice were left untreated, treated with 2 mg of DMPA, or treated with DMPA and 5 days later infected with 1000 FFU of ZIKV intravaginally. All mice were harvested 6 days after ZIKV infection (11 days after DMPA treatment). Cells were isolated from spleens, lymph node, and lower female reproductive tract, and analyzed by flow cytometry. Total cell counts were calculated for T cells, NK cells, B cells, DCs, eosinophils, monocytes, and neutrophils for spleen (**A**), iliac lymph node (**B**) or lower female reproductive tract (**C**). Data represent 4 (Naive and DMPA) or 7 (ZIKV) mice per group, combined from 2 independent experiments. Naive and ZIKV-infected groups were compared to DMPA-treated by two-way ANOVA corrected for multiple comparisons (*, P<0.05; ****, P<0.00001).

Altogether, our data show that although wild-type mice generally do not support ZIKV replication, the vagina is a unique site that supports the replication of ZIKV as well as other flaviviruses. The ability of ZIKV to replicate in the vagina of wild-type mice requires a high progesterone state (pregnancy or DMPA treatment) but the mechanism by which progesterone promotes ZIKV vaginal infection remains unclear.

## DISCUSSION

The emergence of ZIKV in Latin America in 2015-2016 not only revealed new severe disease manifestations but also confirmed a prior report of sexual transmission as an additional mode of transmission for ZIKV, making ZIKV the first arbovirus demonstrated to spread between humans through sexual contact (1, 2). Although most ZIKV cases are presumed to be due to transmission via mosquitoes, it is difficult to estimate to the extent to which sexual transmission contributes to ZIKV transmission in areas with frequent and concurrent mosquito-borne transmission. A retrospective study of ZIKV serology in Brazil found that cohabitating with a ZIKV seropositive sexual partner was associated with a 4-fold greater risk of also being seropositive compared to cohabitating with a ZIKV-seronegative partner whereas cohabitating with a ZIKV-seropositive non-sexual partner was associated with less than a 2-fold greater risk, supporting a role for sexual transmission even in areas with mosquito-borne transmission (28). Sexual transmission may thus have contributed to the high force of infection of ZIKV in this epidemic even in Latin America where any ZIKV cases were presumed to have been acquired via mosquito.

Sexual transmission among humans appears to be an unusual property of ZIKV compared to other flaviviruses, although the incidence and epidemiology of most flaviviruses precludes certainty about the absence of sexual transmission. The best evidence that ZIKV is sexually transmitted is travel-associated cases in the United States, Europe, and elsewhere, wherein women without mosquito exposure became infected after their male partners returned from ZIKV-endemic areas (2, 29-32). Of 5399 travel-associated ZIKV cases in the US 2015-2017, 52 resulted in confirmed transmission to a sexual partner (33, 34). Though this represents only 1% of ZIKV cases in the US resulting in forward sexual transmission, this is likely an underestimate of the rate at which ZIKV-infected men transmit to their partners, since ∼80% of ZIKV infections are asymptomatic and screening has been focused on symptomatic women with travel-related exposure. In contrast, DENV is the most prevalent human flavivirus infection, with an estimated >100 million infections worldwide annually (35) but there have been only two recently-described cases of DENV sexual transmission (36, 37) despite tens of thousands of travel-associated DENV cases over the past >40 years (38-45). Our data in mice suggest that the vagina may be a permissive site for replication of other flaviviruses, as we observed replication of other flaviviruses (SPOV and USUV) that do not generally replicate in wild-type mice (21, 22). Since human infections with those flaviviruses are rare (46, 47), it is not known whether they may share with ZIKV the ability to spread through sexual transmission. It may be that there exists a subset of flaviviruses capable of sexual transmission that have not yet been observed because of the lack of a large enough outbreak for that to be detected. Sexual transmission would also require these viruses to have tropism for the male reproductive tract as well as secretion into semen. Interestingly, SPOV has been observed in semen in mice and to cause fetal pathology in mice, though it has reduced tropism for the male reproductive tract compared to ZIKV (48, 49).

ZIKV pathogenesis often is modeled in *Ifnar1*^*-/-*^ mice to produce robust disseminated infection, including via vaginal inoculation. Though others previously have observed productive ZIKV vaginal infection in wild-type mice (8, 14, 50-52), these studies did not specifically investigate the mechanisms that make the vagina an unusually susceptible site for ZIKV replication in wild-type mice. We found that ZIKV replicates efficiently in the vagina of wild-type mice as measured by viral RNA detectable in vaginal washes and cervix. Remarkably, wild-type mice not only supported ZIKV replication in the vagina but they also sustained equivalent viral loads in the vagina compared to *Ifnar1*^*-/-*^ mice throughout the course of infection. However, only *Ifnar1*^*-/-*^ mice supported systemic infection. These data suggest different roles for IFN-αβ in controlling local ZIKV replication in the vagina versus the disseminated infection. We did not find increased ZIKV replication in the vagina in mice lacking the IFN-λ receptor, contrasting with a prior study reporting that IFN-λ plays a protective role against ZIKV infection in the female reproductive tract (15). The design of the previous study differed from ours in several respects, including using ovariectomized mice supplemented with hormones, treatment with an IFNAR1-blocking antibody, and use of a mouse-adapted ZIKV strain, suggesting that any protective effect of IFN-λ against vaginal ZIKV infection may be context specific.

Importantly, we found that a high progesterone state confers susceptibility to vaginal ZIKV infection in both wild-type and *Ifnar1*^*-/-*^ mice, including high progesterone induced by pregnancy. It is not clear to what extent sex hormones modulate susceptibility to ZIKV infection in humans, though there is precedent for increased HIV susceptibility following progesterone treatment (53). Likewise, progesterone increases susceptibility to HSV in mice (6, 54). The fact that pregnancy in mice causes susceptibility to vaginal ZIKV infection could be important because the most significant outcome of ZIKV infection is congenital infection after either mosquito-borne or sexual transmission (55). The ability of ZIKV to spread sexually creates the potential for congenital infection via an ascending transvaginal route, which would require the virus to cross distinct anatomic and immunologic barriers compared to hematogenous transplacental transmission. It is not known whether an alternative route of congenital infection would be associated with distinct risks and outcomes to the developing fetus. Studies in non-human primates suggest that ZIKV can spread to placenta and fetus following intravaginal inoculation, but the animals in these studies also developed viremia so the route by which the virus spread to the placenta and fetus is uncertain (56, 57).

We found that most ZIKV-infected cells in the vagina were epithelial cells and that there did not appear to be a pronounced immune infiltrate present near sites of infection. The observation that ZIKV infects vaginal epithelial has been reported in mice with impaired IFN-αβ signaling (15). The fact that epithelial cells appear to be the cells primarily infected in vaginal tissue is notable because ZIKV has particular tropism for myeloid cells in systemic infection (26, 27). These data suggest a role for vaginal epithelial cells as mediators of host protection at this site of infection. As pregnant wild-type mice also did not exhibit ascending infection or congenital infection, understanding the mechanisms by which epithelial cells and other cell types restrict ZIKV spread will be important for understanding the risks of sexually transmitted ZIKV in the context of congenital infection.

It previously has been reported that the LFRT expresses lower levels of viral RNA pattern recognition receptors than UFRT, though this expression pattern was not affected by DMPA treatment (8). Accordingly, we found that DMPA did not inhibit baseline or induced expression of *Ifit1*, an antiviral ISG, either in the vagina or the spleen in response to pI:C. Our results suggest that DMPA does not induce a global downregulation of ISG expression that would promote viral infection.

The mechanism by which progesterone confers susceptibility to vaginal ZIKV infection in wild-type mice remains unclear. High progesterone states such as DMPA treatment and diestrus are associated with a thinner vaginal epithelium (7, 54, 58), but we found that vaginal abrasion was not sufficient to permit ZIKV infection in the absence of DMPA treatment, so a compromised epithelial barrier is unlikely to be the primary mechanism by which the vagina becomes susceptible to ZIKV infection. The vaginal epithelium becomes more permeable to leukocytes and microbiota following administration of exogenous progesterone, neutrophil abundance in the vagina increases during diestrus, and progesterone can skew the immune response away from a Th1 towards a Th2 response (54, 59, 60). However, we did not observe a significant change in leukocyte populations systemically or in vaginal tissue after DMPA treatment. Although DMPA potentially could affect specific leukocyte subsets not analyzed here, or affect activation states independently of cell numbers, our results suggest that DMPA-induced susceptibility to vaginal ZIKV infection does not result from a dramatic change in the immune milieu of the vagina. The lack of immune cell infiltrate after DMPA treatment is consistent with prior observations that sex hormones alone do not modulate large changes in immune cell profiles within the LFRT in the absence of infection (61).

Altogether, our results demonstrate that the vagina is an unusually permissive site for ZIKV replication in wild-type mice, but this susceptibility is dependent upon a high-progesterone state, even in immunocompromised mice. The mechanism by which progesterone confers ZIKV susceptibility remains unclear but could include structural changes to the vaginal lumen or epithelial barrier, local or systemic immunomodulatory effects, or direct effects on viral replication in epithelial cells. DMPA treatment is a key component of mouse vaginal infection models for other pathogens, such as HSV and *Chlamydia*, but the mechanisms by which DMPA increases susceptibility to those pathogens also remain poorly defined. Thus, understanding how progesterone mediates susceptibility to ZIKV vaginal infection may provide insights into host mechanisms that influence susceptibility to diverse sexually transmitted pathogens.

## MATERIALS & METHODS

### Cells and viruses

Vero cells were maintained in Dulbecco’s modified Eagle Media (DMEM) supplemented with 5% heat-inactivated fetal bovine serum (FBS) and L-glutamine at 37°C with 5% CO_2_. ZIKV strain H/PF/2013 was obtained from the U.S. Centers for Disease Control and Prevention (62). SPOV strain SA AR 94 and USUV SA AR 1776 were obtained from the World Reference Center for Emerging Viruses and Arboviruses (63, 64). DENV3 WHO reference strain (CH54389) was obtained from Dr. Aravinda de Silva (UNC), RUBV strain M33 from Dr. Michael Rossman (Purdue University) (65) and liver homogenate from HAV infected mice from Dr. Stanley Lemon (UNC) (24).

Virus stocks were grown in Vero cells in DMEM supplemented with 2% FBS and HEPES and titered by focus forming assay (FFA) (66). Virus was serially diluted in duplicate in DMEM supplemented with 2% FBS and HEPES and added to confluent Vero cells in 96 well plates for 1-3 hours at 37°C with 5% CO_2_ before being overlaid with 1% methylcellulose in minimum essential Eagle medium (MEM) supplemented with 2% FBS, HEPES, and penicillin and streptomycin. Cells were then incubated for 40-45 hours at 37°C with 5% CO_2_ before being fixed with 2% paraformaldehyde for 1 hour at room temperature. Cells were then rinsed off with 0.05% Tween-20 in PBS and then incubated for 2 hours at room temperature or overnight at 4°C with 1µg/ml of the flavivirus cross-reactive antibody mE60 (67) in 0.1% saponin and 0.1% bovine serum albumin to permeabilize cells. Following another rinse, cells were then incubated in a 1:5000 dilution of a horseradish peroxidase (HRP) conjugated goat anti-mouse IgG (Sigma). Titration of RUBV was performed similarly but with a polyclonal anti-RUBV goat IgG at 1:4000 (LifeSpan BioSciences, LS-C103273) and a HRP conjugated anti-goat IgG at 1:5000 (Sigma). Color was developed for 30 minutes in TrueBlue substrate (KPL). Foci were quantified using a CTL Immunospot.

UV-inactivated ZIKV was generated by placing 0.2mL ZIKV H/PF/2013 at 1 × 10^6^ FFU/mL in a petri dish and exposing to UV light at 0.9999 J/cm^2^ in an HL-2000 HybriLinker (UVP Laboratory Products) for 10 minutes at room temperature. Mock-inactivated ZIKV was generated similarly but placed under light in a tissue culture hood instead of UV light. Inactivation was confirmed by amplifying UV- and mock-treated virus stocks on Vero cells for 4 days and then titering by FFA.

### Mouse infections

All mouse husbandry and experiments were performed with approval of the University of North Carolina at Chapel Hill’s Institutional Animal Care and Use Committee. All mice were on a C57BL/6J background. *Ifnar1*^*-/-*^ mice were all bred in-house and wild-type mice were either bred on site or purchased from The Jackson Laboratory. Unless otherwise indicated, 5-10 week old female mice were subcutaneously injected with 2mg depot medroxyprogesterone acetate (DMPA) obtained via the UNC pharmacy, diluted in 100µl of PBS. Five days later, mice were challenged with 1000 FFU of virus in 5µl via vaginal instillation or 50µl via footpad. Vaginal abrasion was accomplished by scrubbing the vagina of anesthetized mice with interdental brushes (GUM Proxabrush Go-Betweens tight-sized cleaners) a total of 10 combined full rotations and insertions as previously described (68).

Vaginal washes were collected in a total of 100 µl by twice pipetting 50 µl of PBS with 0.4x protease inhibitor (cOmplete, EDTA-free) into the vagina and collecting immediately, every 2 days after infection. Blood was collected into serum blood collection tubes (BD) days 2 and 6 after infection via submandibular bleed with a 5 mm Goldenrod lancet or via terminal bleed cardiac puncture. Serum was separated at 8000 rpm for 5 minutes. Tissues were collected from mice after euthanasia by isoflurane overdose, cardiac bleed, and perfusion with 5-10 mL of PBS. Tissues, vaginal washes, and serum were stored at -80°C until RNA extraction.

For experiments investigating the responsiveness of tissues to immunogenic RNA, we first treated 5-6 week old mice with either PBS or 2mg DMPA subcutaneously. Four days later, mice were treated with 50 µg polyinosinic:polycytidylic acid (poly(I:C)), low molecular weight (Invivogen, TLRL-Picw) either intraperitoneally in 100µL or intravaginally in 20µL.

### Generation of IFN-λ receptor knock out mice

Mice with a floxed allele of the IFN-λ receptor (*Ifnlr1*^f/f^) were received from Dr. Herbert Virgin (Washington University in St. Louis). *Ifnlr1*^f/f^ mice were crossed with mice expressing Cre recombinase under the β-actin promoter (Jackson Labs # 019099, obtained from Dr. Jenny Ting, UNC) to generate *Ifnlr1*^f/f^ mice with ubiquitous Cre recombinase expression from a hemizygous Cre allele (resulting in *Ifnlr1*^-/-^). These mice were then crossed with *Ifnlr1*^f/f^ mice to generate litters in which 50% of pups lacked IFN-λ signaling (*Ifnlr1*^-/-^, Cre+) and 50% retained it (*Ifnrl1*^+/-^, Cre-). Vaginal infection experiments were conducted in a blinded manner, as genotyping for *Cre* and *Ifnlr1* was performed after the experiment was completed.

### qRT-PCR

RNA from vaginal washes and serum was extracted with the Qiagen viral RNA minikit. RNA from tissues was extracted with the Qiagen RNeasy minikit after homogenization in a MagNA Lyser instrument (Roche Life Science) with zirconia beads (BioSpec) in 600µl PBS followed by incubation at room temperature for 10 minutes in an equal volume RLT buffer for lysis. Viral genomes were quantified by Taqman one-step qRT-PCR on a CFX96 Touch real-time PCR detection system (BioRad) and were reported on a log_10_ scale measured against standard curves from either a ZIKV A-plasmid as previously described (69), or from 400 bp gBlock double stranded DNA fragments (Integrated DNA Technologies, IDT). ZIKV RNA was quantified as previously published (70) and other viruses with the gBlocks and primers in Tables 1 and 2. To measure the expression of *Ifit1* in each tissue, the difference in Ct values between *Ifit1* and *ActB* as a housekeeping gene was calculated for each tissue sample and plotted as -ΔCt.

**Table 1.**
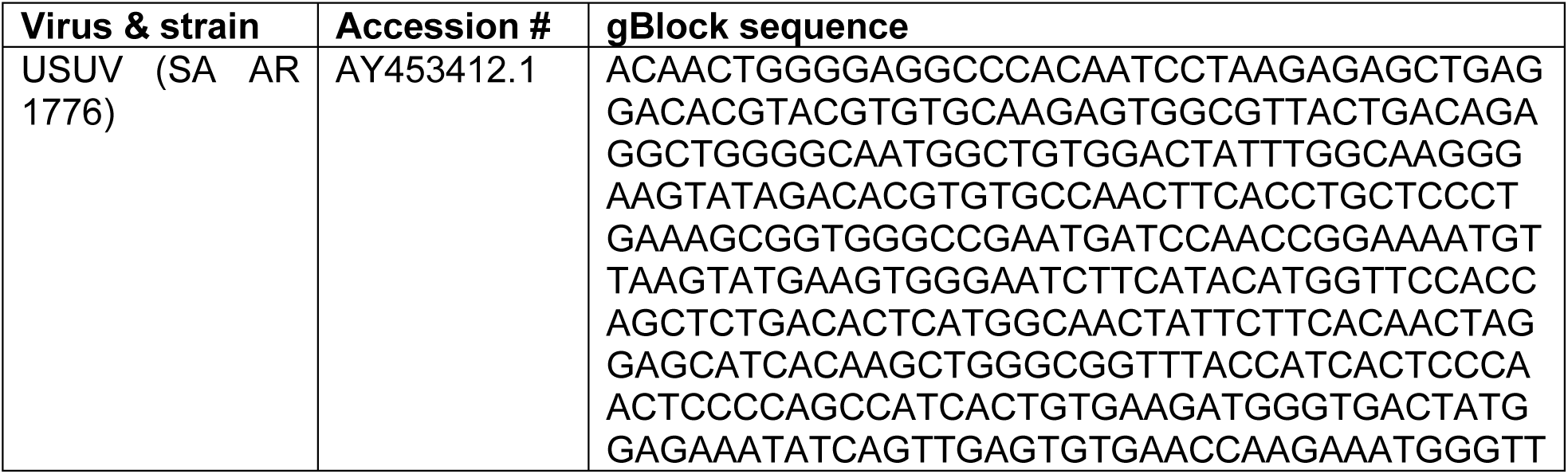
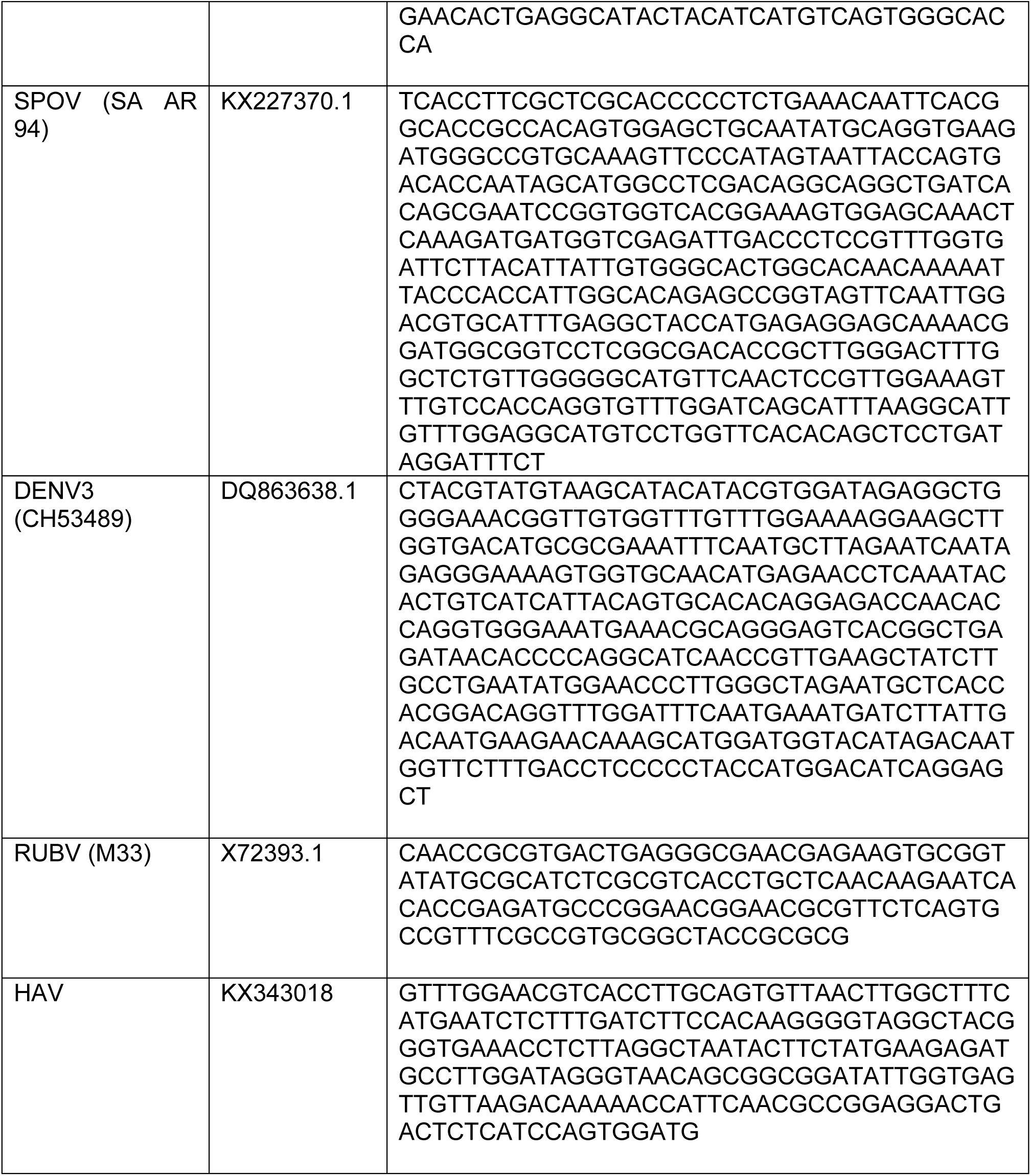
Sequences used for qPCR standard curves.

**Table 2.**
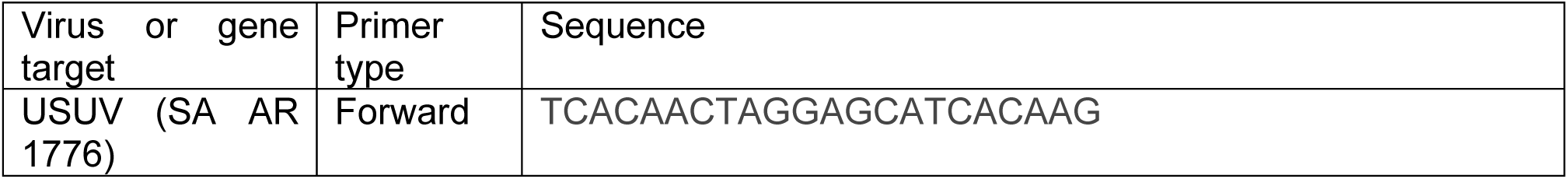
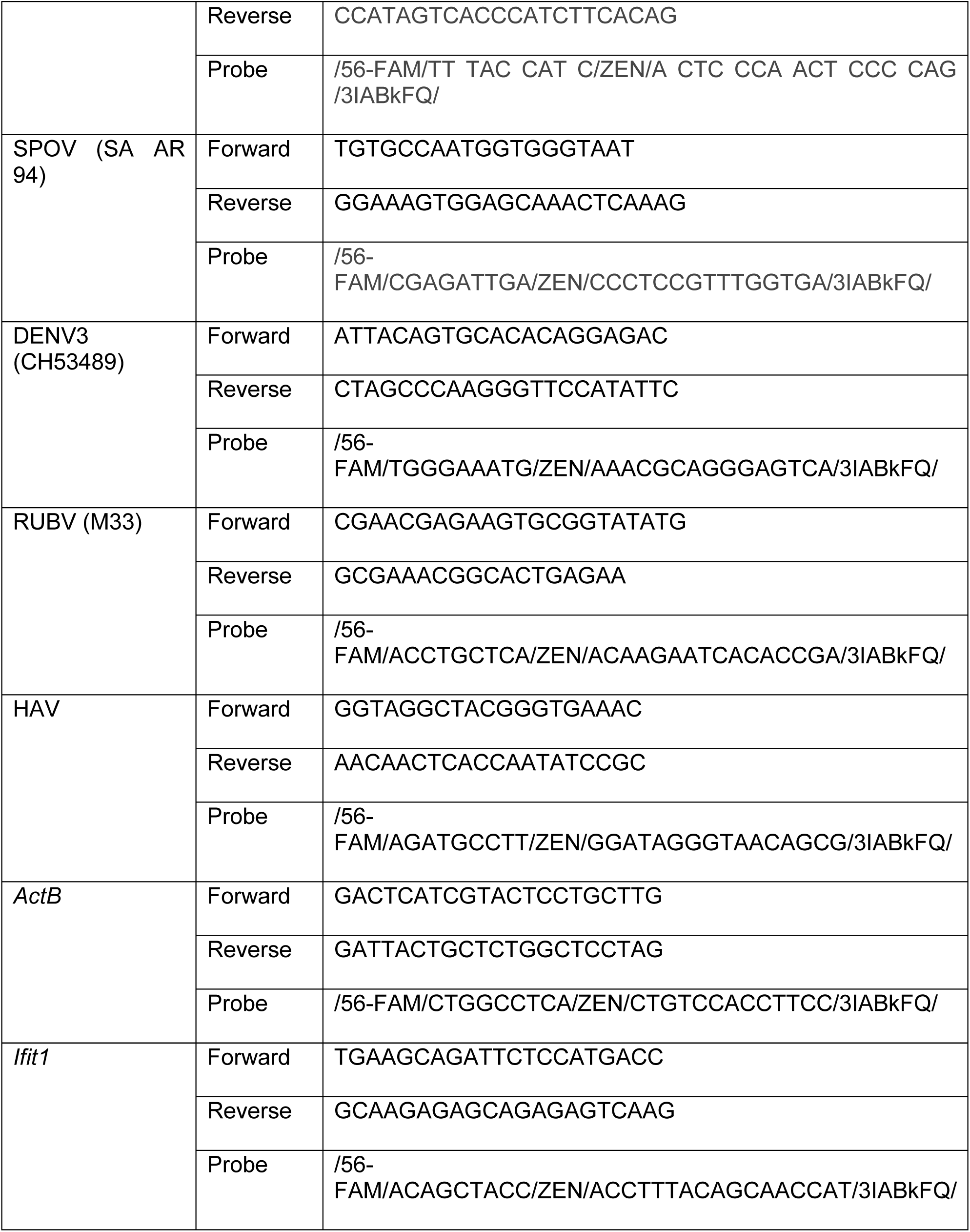
Primer sets used for qRT-PCR.

### In situ hybridization

Tissues were collected from euthanized mice after exsanguination by cardiac puncture and perfusion with 10 mL of PBS followed by 10 mL of 10% neutral buffered formalin (NBF). Tissues were then stored overnight in 1mL of 10% NBF at 4°C before being transferred to PBS at 4°C for longer term storage. Tissues were paraffin embedded and 5µm sections stained with a ZIKV-specific RNA probe (Advanced Cell Diagnostics #467871) and a hematoxylin counter-stain. Positive and negative staining controls for RNA-specific staining were confirmed with probes against peptidyl-prolyl cis-isomerase B (PPIB, #321651) and dihydrodicipicolinate reductase (dapB, #320751) as recommended by the manufacturer. Tissue processing, histology, and RNAscope was performed by the UNC Histology Research Core Facility.

### Flow cytometry

Spleens and iliac lymph nodes (iLNs) were mechanically dissociated and red blood cells were lysed using RBC lysis buffer (0.84% NH4Cl in PBS). Cells were pelleted by centrifugation and resuspend in media (RPMI 1640 with 1% FBS). Cells were filtered through a 70 μm cell strainer to make a single-cell suspension. LFRT tissue was excised, minced with scissors, and digested in HBSS (with Ca^2+^ and Mg^2+^) containing 1 mg/mL Collagenase I and 0.05 mg/mL DNAse I for 60min at 37°C in a shaking incubator. After incubation, 1mL FBS was added to stop digestion and cells were serially filtered through a 40- and 70-μm cell strainer and washed with HBSS (with Ca^2+^ and Mg^2+^). Cells were resuspended in media at a concentration of 1×10^7^ cells/mL for flow cytometric analysis.

Isolated cells were stained in PBS with 1% FBS for 20–30 min in the dark on ice. Fc receptor blockade was performed with anti-CD16/32 mAb prior to surface staining. Dead cells were excluded from analysis using Zombie UV (BioLegend). Cells were fixed in 2% paraformaldehyde, and samples were acquired using an LSRII flow cytometer (BD Biosciences). Data were analyzed using FlowJo software (Tree Star). The following antibodies were used in this study: anti-CD16/32 (clone 2.4G2; BD Biosciences), anti-CD45 AF700 (clone 30-F11; BioLegend), anti-CD3e APC-Fire/750 (clone 17A2; BioLegend), anti-CD19 PE-Cy7 (clone 6D5; BioLegend), anti-NK1.1 PE (clone PK136; BioLegend), anti-CD11b APC (clone M1/70; BioLegend), anti-CD11c BV650 (clone N418; BioLegend), anti-Ly6G FITC (clone IA8; BioLegend), and anti-Ly6C BV605 (clone HK1.4; BioLegend). The following markers were used to identify immune cell populations: T cells (CD45+CD3e+), B cells (CD45+CD19+), NK cells (CD45+NK1.1+), dendritic cells (CD45+CD11c+), neutrophils (CD45+CD11b+Ly6G+), and monocytes (CD45+CD11b+Ly6G-Ly6C+/-).

### Statistical analysis

Statistical tests were performed with Graphpad Prism 9.0. Tests used include unpaired multiple Mann-Whitney analyses with the Holm-Śídák method and two-way ANOVA with matched time points where multiple time points of the same mouse were taken, the Geisser-Greenhouse correction for lack of sphericity, comparison to control cell means, and the Dunnett correction for multiple comparisons.

## ACKNOWLEDGEMENTS

This work was supported by R21 AI144631 (H.M.L.) and start-up funds from UNC Chapel Hill Department of Microbiology & Immunology and the Lineberger Comprehensive Cancer Center. H.M.L. holds an Investigators in the Pathogenesis of Infectious Disease Award from the Burroughs Wellcome Fund. C.A.L. was supported by F31 AI143237; S.J.D. was supported by K12 GM000678. Histology services were provided by the UNC Histology Research Core Facility. The UNC Flow Cytometry Core Facility is supported in part by P30 CA016086 Cancer Center Core Support Grant to the UNC Lineberger Comprehensive Cancer Center.

